# Designing new natural-mimetic phosphatidic acid: a versatile and innovative synthetic strategy for glycerophospholipid research

**DOI:** 10.1101/2025.02.20.639292

**Authors:** Antoine Schlichter, Alexander Wolf, Thomas Ferrand, Aurelien Cocq, Lina Riachy, Steven Vertueux, Brice Beauvais, Marine Courvalet, Paul-Joël Henry, Emeline Tanguy, Louis Gonzales, Rémy Ferlet, Fanny Laguerre, Charles Decraene, Alexia Pellissier, Muriel Sebban, Cyrille Sabot, Lydie Jeandel, Sarah Cianférani, Jean-Marc Strub, Magalie Bénard, Victor Flon, Valérie Peulon-Agasse, Pascal Cardinael, Stéphane Ory, Stéphane Gasman, Pierre-Yves Renard, Maïté Montero-Hadjadje, Nicolas Vitale, Sébastien Balieu

## Abstract

Glycerophospholipids (GPLs) play important roles in cellular compartmentalization and signaling. Among them, phosphatidic acids (PA) exist as many distinct species depending on acyl chain composition, each one potentially displaying unique signaling function. Although the signaling functions of PA have already been demonstrated in multiple cellular processes, the specific roles of individual PA species remain obscure due to a lack of appropriate tools. Indeed, current synthetic PA analogues fail to preserve all the functions of natural PA. To circumvent these limitations, we developed a novel synthetic approach to produce PA analogues without compromising structural integrity of acyl chains. Moreover, addition of a clickable moiety allowed flexible grafting of different molecules to PA analogues for various biological applications. Hence, this innovation also provides powerful tools to investigate specific biological activities of individual PA species, with potential applications in unraveling complex GPL-mediated signaling pathways.

## Introduction

Lipids are essential constituent of organisms, yet their extreme diversity and lack of appropriate tools overcomplicate their studies^1,2^. Glycerophospholipids (GPLs) are a subclass of lipids, constituting the bulk of cellular membranes, but are also able to perform important signaling functions within cells. Coexistence of hundreds of GPLs within biological membranes suggests that each plays major, yet potentially, non-overlapping roles. Furthermore, given their rapid and complex metabolism, establishing the roles of an individual lipid, in a spatial and temporal resolved manner, remains challenging^3^. Among GPLs, phosphatidic acids (PA), composed of a single phosphomonoester headgroup (the smallest headgroup within GPL), have been shown to exhibit structural diversity within cells based on their fatty acyl chains composition^4,5^. PA has attracted considerable attention and has been proposed to play pivotal roles in multiple cellular functions^3,4^. Accordingly dysregulation of its metabolism is linked to numerous diseases including neurodegeneration^6^, intellectual disabilities^7,8^, cancer^9^, fragile X^10^ and Coffrin-Lowry syndromes^11^. Currently available GPL analogues^12^, aiming at understanding their individual role, rely on modified fatty acyl chains (with fluorophore,^13^ photo-crosslinker^14^ or a clickable moiety^15^), or modified headgroups, which is not applicable to PA. However, side chains composition is critical for biological activity of GPLs and altering them impacts their ^16,17,18,19^ functions. To preserve biological properties of GPLs, it is anticipated that both their headgroups and fatty acyl chains^20^ must remain unchanged. Hence, the glycerol backbone might be the most relevant place to introduce chemobiological modifications. This approach was first proposed by adding a clickable azidomethyl moiety on the glycerol core^21^. *In vitro* studies demonstrated that this modification had little effect on PA probe association with protein kinase C (PKC), validating this strategy. However, the proposed synthesis only allowed grafting two identical fatty acyl chains, which does not match the diversity of natural GPLs. To circumvent these limitations, we developed a novel synthetic strategy for clickable azide-based GPL analogues (GPL-N_3_), allowing to keep untouched fatty acyl chain composition and phosphate head group, thus mimic natural GPL species, and first applied this strategy to PA, where phosphate headgroup modification is not applicable. The specificity of those novel PA analogues was validated in chromaffin cells, a model in which we recently demonstrated that different PA species have exclusive non-redundant functions in the secretory pathway and allowed confirmation of previously identified PA-interacting proteins involved in this process but also unveiled dozens of new potential PA binding partners.

## Results and discussion

### Convergent synthesis of GPL-N_3_ and GPL probes

We developed a new specific synthetic route to achieve regio-selective and stereo-specific grafting of fatty acyl chains. This method incorporates a clickable moiety on the glycerol core of GPL structure (Figure 1). Headgroup protection as dimethylphosphoester facilitated purifications steps of various intermediates. To obtain accurate stereochemistry of the *sn-*2 position, synthesis was initiated with dimethyl-L-tartrate. Several transformations allowed us to differentiate the four carbon atoms of the tartric backbone resulting in intermediate **8**, the common building block for all GPL-N_3_.

**Figure 1:**
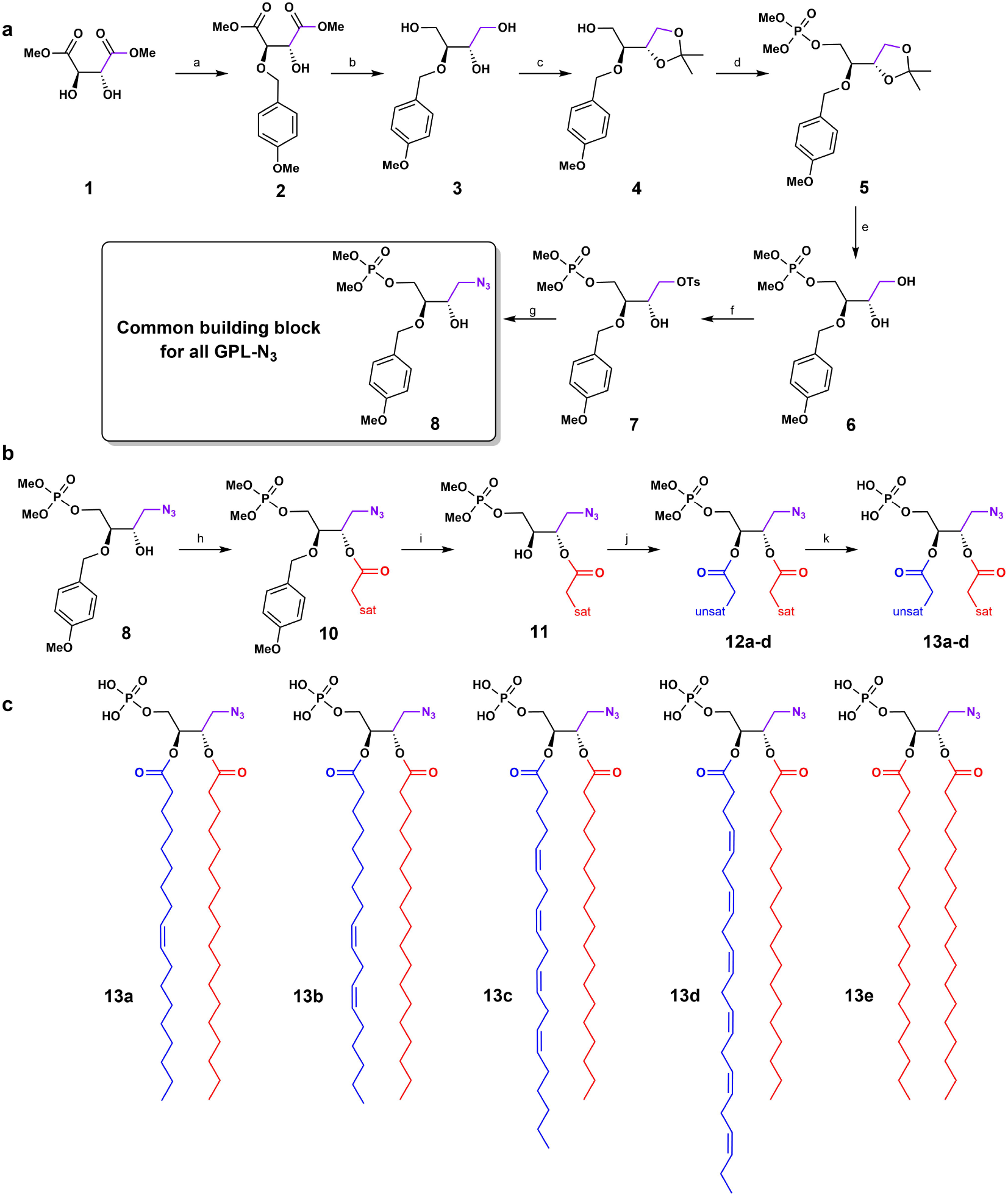
Synthetic route to the common building block 8 leading to PA-N_3_ 13a-d. **a**, formal synthesis of the common building block **8** having all the *sn* positions differentiated. a) PMBBr, Bu_2_SnO, Bu_4_NI, CsF, Toluene, 70%. b) NaBH_4_, *i*-PrOH, 90%. c) 2,2Dimethoxypropane, APTS, acetone, 70%. d) Dimethylchlorophosphate, *t*-BuOK, DCM, 76%. e) HCl, MeOH, 85%. f) TsCl, DIPEA, Bu_2_SnO (0.5% mol.), toluene, 75%. g) 1) NaN_3_, DMF, 2) TMSCHN_2_, MeOH, 65%. **b**, formal synthesis of four different PA-N_3_ **13a-d** from intermediate **8**. h) DCC, DMAP, stearic acid, DCM, 93%. i) DDQ, DCM, PBS buffer (pH= 7.2). j) DCC, DMAP, oleic acid (**12a**) or linoleic acid (**12b**) or arachidonic acid (**12c**) or docosahexaenoic acid (**12d**), 50% (**12a, 12b**), 63% (**12c**) 75% (**12d**) over two steps. k) 2-methylbut-2-ene, TMSBr, CHCl_3_ or 2-methylbut-2-ene, TMSBr, BSTFA, CHCl_3_, 48 to 95%. **c**, structures of the PA-N_3_ (18:0-18:1) **13a**, PA-N_3_ (18:0-18:2) **13b**, PA-N_3_ (18:0-20:4) **13c**, PA-N_3_ (18:0-22:6) **13d** and PA-N_3_ (18:0-18:0) **13e**.

The *sn*-3 position was functionalized with dimethyl phosphate ester, the *sn-*2 was protected with a *p*-methoxylbenzyl (PMB) group, and the *sn-*1 position was let free. The additional carbon on the glycerol core, designated as *sn-*0, was equipped with an azide moiety. Despite high yields of the first synthetic steps (Figure 1a and SI 1.5.1), an unexpected deprotection of the phosphate ester occurred during the nucleophilic substitution of tosylate **7** by sodium azide leading to **9**. Although reaction between lithium azide, and to a lesser extent sodium azide, is well known for cleaving methyl phosphate monoesters^22^, it was expected that tosylate displacement by an azide would be selective over this ester cleavage. However, this was not the case despite modifications in temperature, number of equivalents, azide source, solvents and leaving groups tested. The optimal solution found was to realkylate the deprotected phosphate of **9** using TMSCHN 2^23^ giving **8** in an overall 14% yield in seven steps (Figure 1a). Using this synthetic route, **8** could be obtained at the gram scale.

The second part of the synthetic route afforded four different PA-N_3_ **13a-d** from the common building block **8**, with regio-selective addition of desired fatty acids through sequential acylation/deprotection/acylation steps. The *sn*-1 position was acylated with a saturated fatty acid (stearic acid), leading to ester **10** with good yield thanks to the coupling reagent DCC/DMAP. Subsequent deprotection of PMB moiety was carefully monitored in mild conditions by ^31^P NMR until completion, immediately followed, without purification, by a second acylation reaction with an unsaturated fatty acid with the same coupling agent DCC/DMAP. This reaction converted resulting *sn*-2 alcohol into esters **12a-d** as quickly as possible avoiding any significant intramolecular trans-esterification reaction with stearate. Velocity of this side-reaction was studied by NMR with shorter fatty acids chains and was well characterized. Thus, in our reaction conditions (0 °C, 5 h or room temperature, 1 h) we avoided this side-reaction and consequently maintained regioselectivity higher than 95% during grafting of fatty acyl chains (see transesterification studies in SI 1.6). Finally, cleavage of dimethyl phosphate ester of **12a-e** was performed under mild conditions with BSTFA^24,25^ and TMSBr to obtain various PA-N_3_ **13a-d**, (PA-N_3_ (18:0-18:1) **13a**, PA-N_3_ (18:0-18:2) **13b**, PA-N_3_ (18:0-20:4) **13c**, PA-N_3_ (18:0-22:6) **13d**), whereas symmetrical PA-N_3_ (18:0-18:0) **13e** was also obtained through a shorter synthetic pathway described in supplementary materials (Figure 1b-c and SI 1.5.2). This process leading to **13a-d** is amenable to any other fatty acyl chain composition, thus allowing us to accordingly access PA structural diversity.

To produce PA probes from PA-N_3_ **13a-d**, we performed SPAAC ligation reaction to avoid generation of ROS species with copper (I)^26^; even though this reaction is slower than CuAAC^27^, SPAAC has proved to be amenable to *in vivo* chemistry within life cells. The chosen cyclooctyne SPAAC partners were obtained from DIBAC derivatives offering a good compromise between stability and reactivity. The DIBAC-C_6_-Amine synthesis^28^ was optimized to minimize side products formation and improve reaction repeatability, especially for the dibromation/elimination sequence leading to the cyclooctyne ring (SI 1.5.3). Thereafter a coupling reaction with activated carboxylic acid derivatives was performed affording photocrosslinker (PCL)-DDA1 **18** and DIBAC-C_6_-ATTO 647N **19** or SNAr reaction to get DIBAC-C_6_NBD **20** (Figure 2a). These chemobiological tools were first developed with PA-N_3_ (18:0-18:1) **13a and PA-N**_3_ (18:0-22:6) **13d** due to their non-overlapping functions in neurosecretion^29^. SPAAC reactions led to six new PA probes bearing either diazirines^30^ as photocrosslinkers with PA-DDA1 (18:0-18:1) **21a** and PA-DDA1 (18:0-22:6) **21d** or either fluorophore with PANBD (18:0-18:1) **22a**, PA-NBD (18:0-22:6) **22d**, PA-ATTO 647N (18:0-18:1) **23a** and PA-ATTO 647N (18:0-22:6) **23d** (Figure 2b).

**Figure 2:**
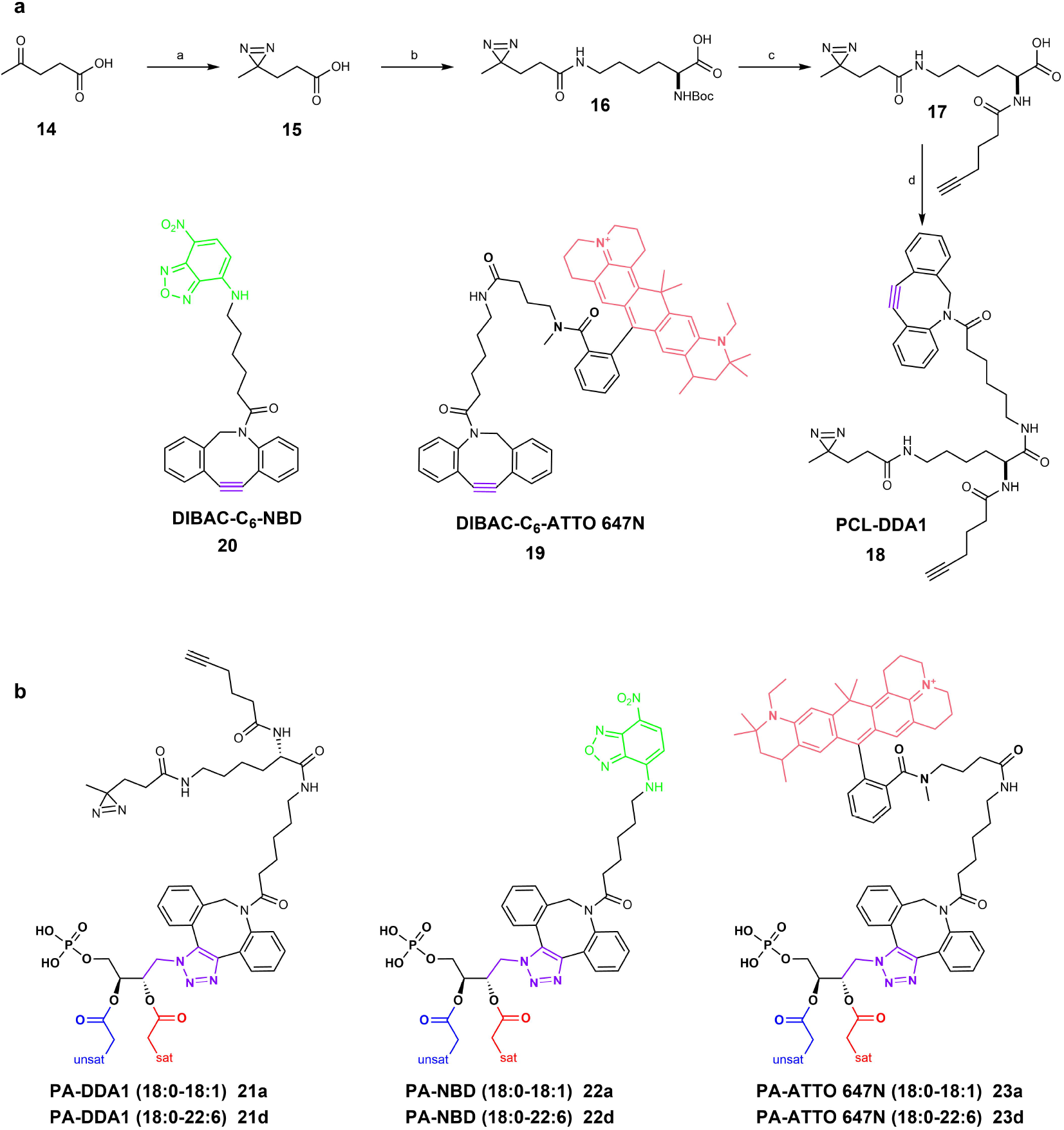
Synthesis of the PA analogues. **a**, formal synthesis of the PCL-DDA1 **18** and structure of the synthesized clickable fluorophore DIBAC-C6-ATTO 647N **19** and DIBAC-C_6_NBD 20. a) 1) NH_3_, MeOH, *t-*BuOCl, 2) *t-*BuOCl b) 1) NHS, EDCI, DMF, 2) Boc-Lys-OH, DIPEA, DMF, 80%. c) 1) TFA, DCM, 2) Pent-5-yn-*N*-hydroxysuccinimide ester, 50% over two steps. d) 1) NHS, EDCI, THF, 2) DIBAC-NH_2_, DIPEA, rt, 20% over two steps. **b**, structures of the PA probes **21-23** obtained by SPAAC click-chemistry between cyclooctynes **18**-**20** and PAN3 13a and 13d.

### Biocompatibility and membrane incorporation of synthetic mono-and poly-unsaturated PA analogues in neurosecretory cells

Primary chromaffin cell live-imaging revealed that PA-ATTO 647N (18:0-18:1) **23a** and PAATTO 647N (18:0-22:6) **23d** analogues rapidly inserted into cellular membranes and accessed intracellular compartments without visible alteration (Figure 3a-b and video 1-3). PA-ATTO 647N (18:0-22:6) **23d** displayed faster access to the plasma membrane and accumulated more effectively in internal membrane compartments than PA-ATTO 647N (18:0-18:1) **23a**, in agreement with the established cellular gradient of poly-unsaturated GPL, decreasing from intracellular organelles towards the plasma membrane^31^. Validating the rapid access of PA analogues to the inner leaflet of the plasma membrane, the PA sensor Spo20p-GFP^32^ was recruited to the plasma membrane within minutes of incubation of PA analogues **13a, 13d, 21a** and **21d** (Figure 3c-d). These data confirm integrity and stability of synthetic PAs *in cellulo*, but also their ability to interact with a known PA binding protein, and highlight their potential utility to study membrane dynamics and signaling processes in live cells.

**Figure 3:**
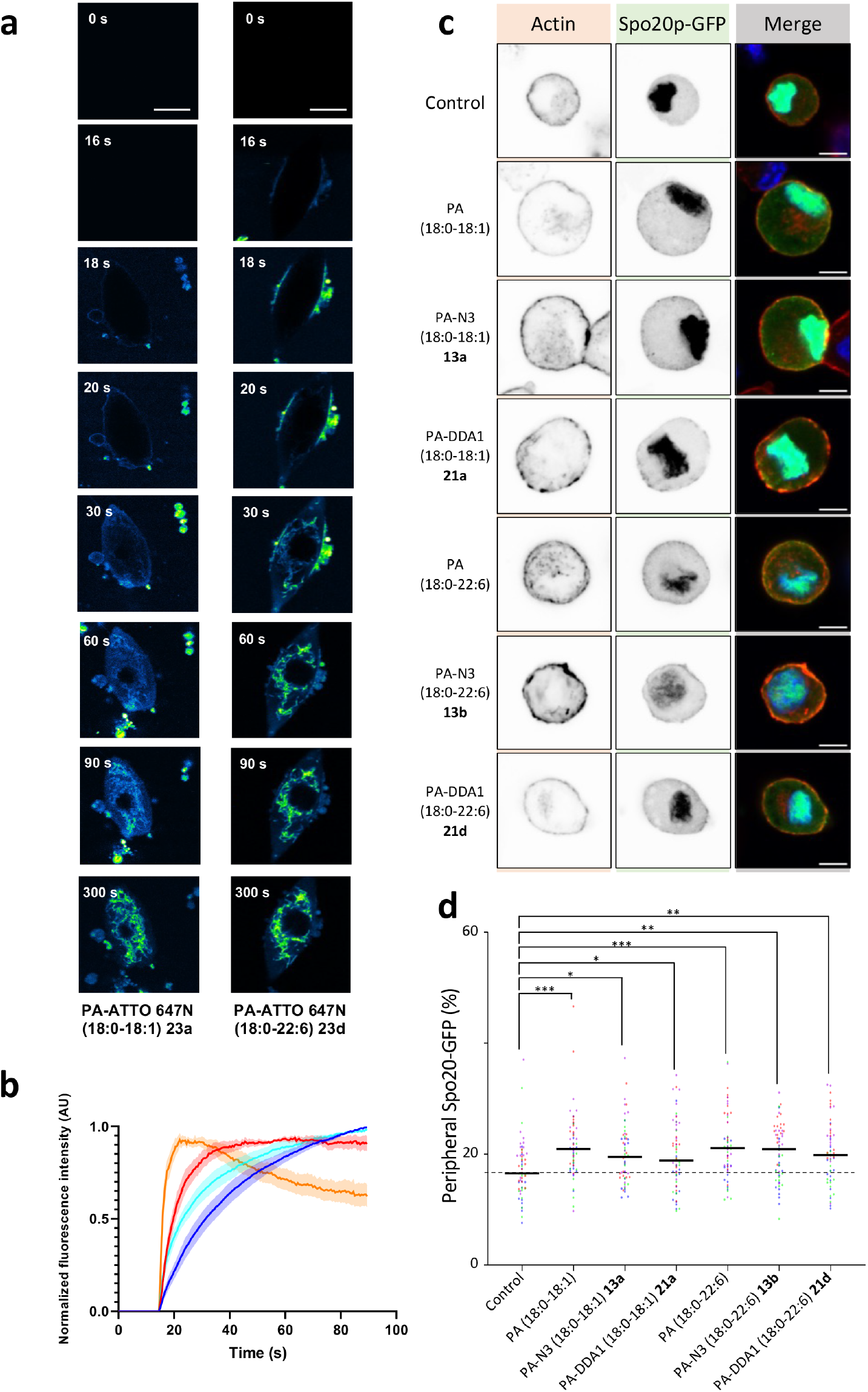
Membrane incorporation of synthetic PA analogues. **a**, Live-cell imaging showing the incorporation kinetics of 3 μM fluorogenic PA-ATTO 647N (18:0-18:1) **23a** or PAATTO 647N (18:0-22:6) **23d** on bovine chromaffin cells. Images have been false colored with ImageJ’s Green Fire Blue LUT (intensity: blue<green<yellow). Biomolecules have been added on living cells 15 s after the beginning of recording (arrow on graph). Images were acquired at a speed of 2 frames per second for 90 s. Scale bar, 10 μm. **b**, Mean fluorescence intensity was analyzed in the whole cell (dark blue line: **23a**; light blue line: **23d**) and in a peripheric area of 500 nm width (red line: **23a**; orange line: **23d**). AU, arbitrary units. The maximum fluorescence intensity was normalized to 1. Analysis was performed on 6 cells from 4 independent cell cultures. Data are expressed as mean ± SEM.; **c**, Recruitment of the PA sensor Spo20p-GFP after addition of 100 μM natural PA or synthetic PA analogues. Bovine chromaffin cells overexpressing the PA sensor Spo20p-GFP were incubated with the mentioned natural PAs or synthetic PA analogues for 15 min prior to fixation. Representative confocal images are shown, with phalloidin-Atto647N (Actin) and Spo20p-GFP (PA sensor) channel displayed individually as an inverted grayscale image and merged (Blue: nucleus, red: actin and green: Spo20p-GFP). Scale bar, 5 μm. **d**, quantification of plasma membrane recruitment of PA sensor Spo20p-GFP. Data are represented as median, and each point is a measure from an individual cell. Points are colored coded (red, blue, green or magenta) according to experimental repeats. Basal recruitment of Spo20p-GFP is indicated by the dotted line. *: p<0.05, **: p<0.01, ***: p<0.001, Kruskall-Wallis followed by Dunn’s multiple comparisons test vs control (N=60 per condition, 15 cells from 4 experimental repeats performed on 2 independent cultures).

### Functional validation of synthetic PA analogues in the secretory process

To functionally validate these new synthetic PA analogues, we tested their ability to participate in membrane remodeling: interaction between chromogranin A (CgA) glycoprotein, and PA (18:0-18:1) being able to induce such membrane remodeling. In presence of CgA-AF633, giant unilamellar vesicles (GUVs) containing PA-NBD (18:0-18:1) **22a** exhibited significant budding events in agreement with what was previously observed^33^. As expected, fluorescence microscopy images showed colocalization of both partners. In contrast, CgA-AF633 failed to induce the budding of GUVs containing commercially available PA (18:1-6:0-NBD) bearing a modification on fatty acyl chains probably because the modified acyl chain loop up to the surface of the vesicle^34^ (Figure 4a). Quantitative analysis confirmed that in presence of CgA we observed significantly more budding GUVs when they were formed with PA-NBD (18:0-18:1) **22a** than with PA (18:1-6:0-NBD) (Figure 4b), demonstrating that *(i)* CgA interacts effectively with synthetic PA-NBD (18:0-18:1) analogue **22a**, and *(ii)* this interaction is functional, inducing membrane remodeling and vesicle budding, mimicking cellular properties of native PA (18:0-18:1) in Golgi and secretory granule membranes. Conversely, commercial PA (18:1-6:0-NBD) failed to replicate those effects.

**Figure 4:**
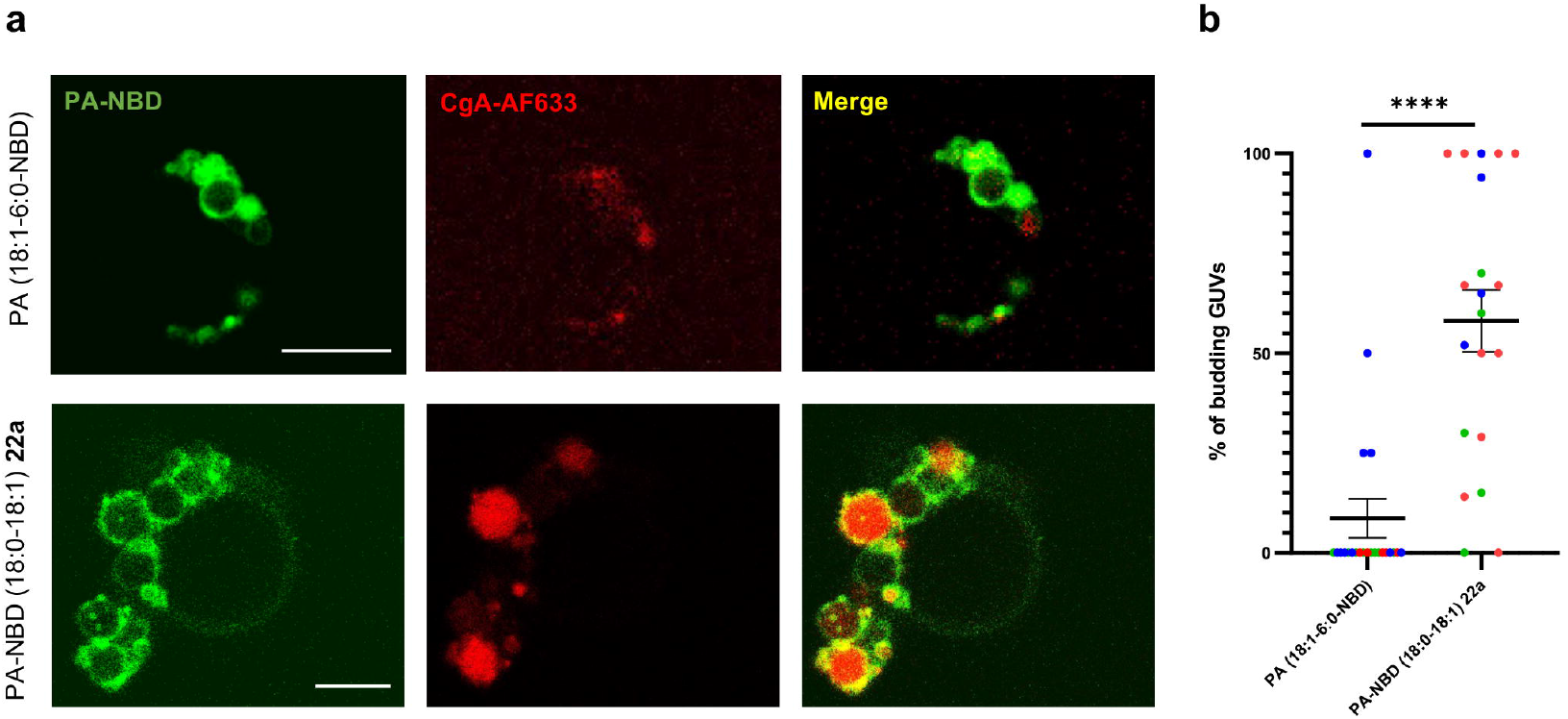
Liposome-based evaluation of synthetic fluorescent PA-NBD analogues. **a**, representative pictures of GUVs composed of 96% DOPC and 4% PA (18:1-6:0-NBD) or 4% PA-NBD (18:0-18:1) **22a**, in the presence of CgA-AF633. Scale bar, 10 μm. **b**, quantification of budding GUVs composed of 96% DOPC and 4% PA (18:1-6:0-NBD) or 4% PA-NBD (18:018:1) **22a** with CgA-AF633. Each point represents the result of GUV analysis from three independent experiments (20 and 22 images for each type of PA-NBD containing-GUVs, respectively). Points are colored coded (red, blue or green) according to experimental repeats. ****: p<0.0001, Mann Whitney test.

PA synthesized by phospholipase D (PLD) promotes secretion from various cell types including neurons, endocrine and exocrine cells^35,36,37^. For instance, we previously reported that knock out, knock-down, or pharmacological inhibition of phospholipase D1 (PLD1) impaired catecholamine release from chromaffin cells by altering multiple steps of the ^29,38^ neurosecretory process. Interestingly, extracellular provision of mono-unsaturated PA (18:0-18:1) restored the number of exocytotic events, whereas extracellular provision of polyunsaturated PA (18:0-22:6) regulated the kinetics of individual events. Hence, to assess whether synthetic PAs recapitulate these specific biological effects, we used DβH labeling of exocytotic spots combined with amperometric measurements as reporters of the secretory activity of chromaffin cells. Potent inhibition of DβH staining induced by the PLD1 inhibitor CAY93 was efficiently rescued by synthetic PA (18:0-18:1) whether they harbored a N_3_-linker **13a**, a fluorescent NBD group **22a**, or a PCL DDA1 moiety **21a** (Figure 5a-b). Importantly, synthetic PA-N_3_ (18:0-22:6) **13d** and its derived DDA1 probe **21d** failed to rescue DβH labeling back to the control condition, in agreement with data obtained with natural PA (18:0-22:6). Moreover, addition of acyl chain modified PA (16:0-6:0-NBD) was also unable to rescue the effects of PLD1 inhibition (Figure 5a-b). Furthermore, electrochemical recording of catecholamine secretion on PLD1 inhibited chromaffin cells enabled us to explore the specific contribution of PA (18:0-22:6) and PA-DDA1 (18:0-22:6) **21d**. CAY93 treatment reduced the number of measured exocytotic events, an effect that was not rescued by addition of natural and synthetic poly-unsaturated PA species (Figure 5c-f and h). However, CAY93 also slowed down the speed of individual fusion events (Figure 5g and i) and increased the duration of individual fusion pores (Figure 5j). Both those effects were effectively rescued by PA (18:0-22:6) and PA-DDA1 (18:0-22:6) **21d**. Hence, the synthetic PA analogues developed recapitulate exclusive functions of natural mono-and poly-unsaturated forms of PA during neuroendocrine secretion, a capability not observed with PA (18:0-6:0-NBD). These data strongly support the idea that modifications targeting the glycerol backbone of PA do not alter functional properties of these PA analogues within cells.

**Figure 5:**
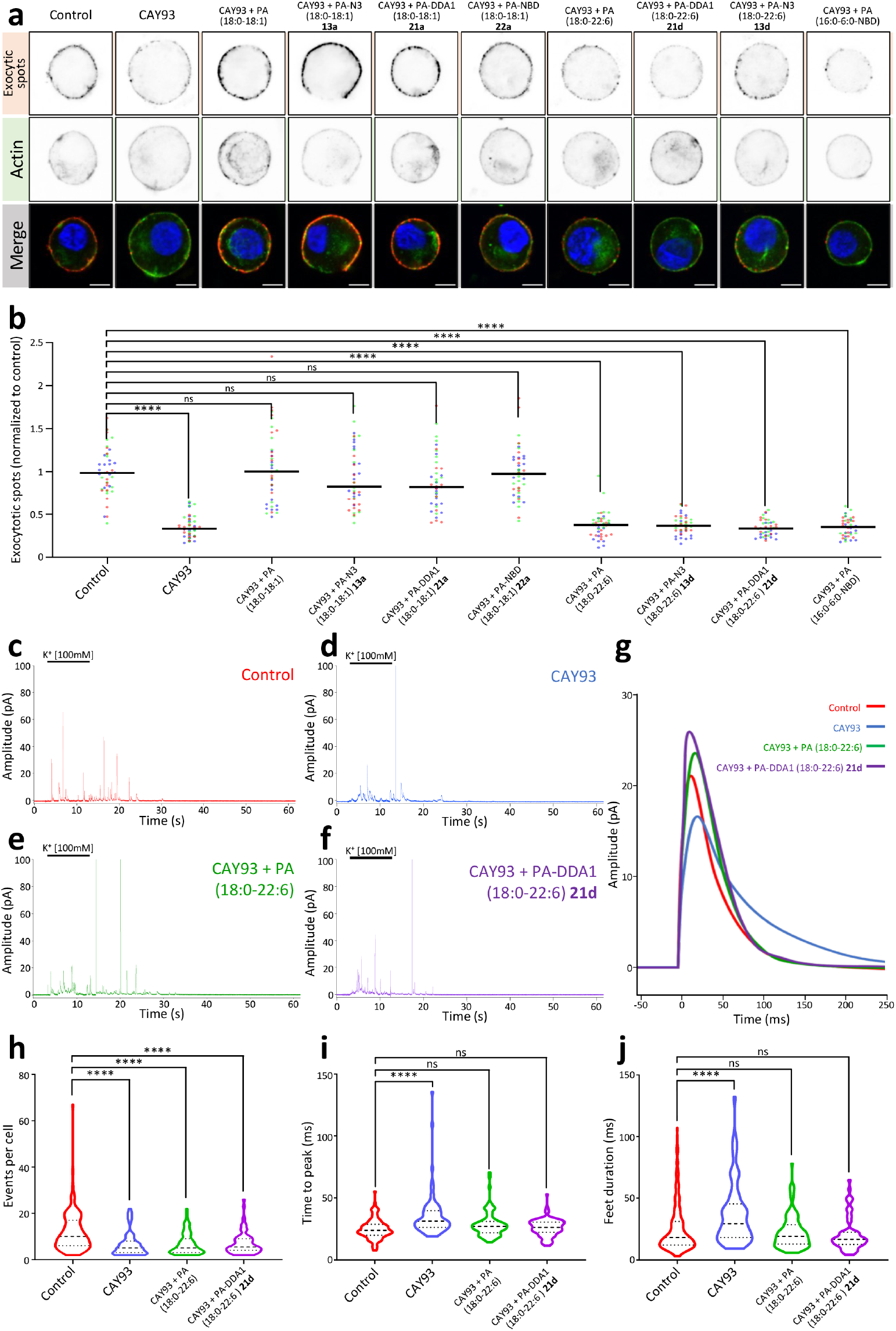
Synthetic PA analogues fully recapitulate the functions of natural PA in neuroendocrine secretion. **a**, representative confocal images of stimulated bovine chromaffin cells after exocytotic spots staining. PLD1 activity was inhibited with 50 nM CAY93, and cells were incubated with natural PAs and synthetic PA analogues. Exocytotic spot (DβH staining) and actin network (phalloidin-Atto647N staining) are displayed individually as an inverted grayscale image and merged (Blue: nucleus, red: exocytotic spots and green: actin). Scale bar, 5 μm; **b**, exocytotic spots quantification (D□H staining). Peripheral exocytotic spot intensity has been normalized with the mean control condition of the individual experimental repeat. Data are represented as median, and each point is a measure from an individual cell. Symbols are colored coded (red, blue, green or magenta) according to experimental repeats. ns: non-significant, ****: p<0.0001, Kruskall-Wallis followed by Dunn’s multiple comparisons test (N=45 per condition, 15 cells from 3 independent cultures). Without PLD1 inhibition PA addition does not affect overall secretion of cells upon stimulation and specific effects of PA (18:0-22:6) were also reproduced in resting condition (Extended figure 1a). Cell area was not affected by any of the treatments (Extended figure 1b). **c-f**, examples of amperometrical traces from cells incubated with or without PLD1 inhibitor (50 nM CAY93) and the mentioned PA species, or PA analogue. Cells were stimulated with a 100 mM K^+^ solution (Black line on each trace). **g**, schematic representation of mean spikes obtained from those recordings. **h**-**j**, violin plots representing respectively the number of exocytotic events, spike time to peak and feet duration of individual spikes (Bars in the plot represent from top to bottom, third quartile, median and first quartile). ns: non-significant, ****: p<0.0001, Kruskall-Wallis followed by Dunn’s multiple comparisons test vs control (N>82 per condition from 3 independent cultures), other parameters were also quantified and yielded similar results (Extended figure 2).

### Characterization of the PA interactome in neurosecretory cells using synthetic PADDA1

To further assess functional differences between mono-and polyunsaturated PA, we performed photo-crosslinking experiments to map the various PA interacting proteins landscape during neurosecretion. Chromaffin cells were incubated with PA-DDA1 (18:0-18:1) **21a** or PA-DDA1 (18:0-22:6) **21d** and illuminated to crosslink PA interactants in resting, stimulated, or post-stimulation states or without illumination. Synthetic PAs were then purified from cell lysates using the alkyne moiety of DDA1 by performing a CuAAC reaction with azide tagged beads. Finally, interacting proteins were identified by mass spectrometry following trypsin digestion of the purified samples. A total of 1264 polypeptides identified without illumination were considered as background. After subtracting them, we identified 778 potential direct or indirect PA partners. Validating our strategy, these include proteins previously identified as PA interactants in assays using cell extracts incubated with PA coupled to agarose beads^39,40^ and known PA interactants validated biochemically^41,42,43,44^ (Extended figure 4). Additionally, many novel potential PA partners were discovered. Among these, 286 were specific to PA (18:0-18:1) **21a** and 341 to PA (18:0-22:06) **21d**, whereas 151 were common to both PA species (Figure 6a). Furthermore, some of these interactants were specific to either resting, stimulated, or post-stimulated cells, whereas others were present in two or three conditions (Figure 6a), highlighting the dynamic nature of these interactions.

**Figure 6:**
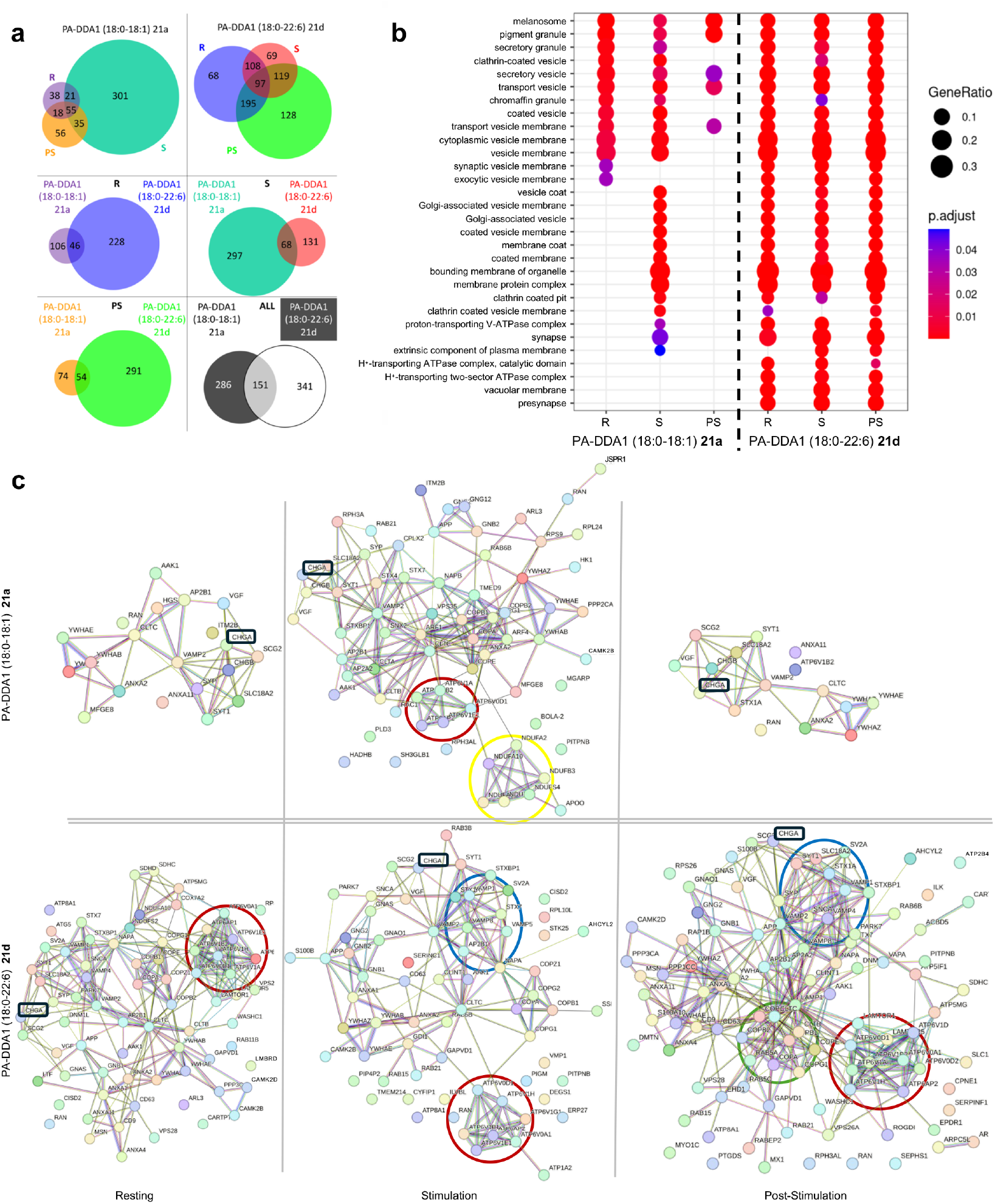
PA interactome in neurosecretory cells. **a**, distribution of different proteins identified in the fishing experiments performed in bovine chromaffin cells for PA-DDA1 (18:018:1) **21a** and PA-DDA1 (18:0-22:6) **21d** in resting condition (R), stimulated condition (S), or resting condition post-stimulation (PS). **b**, cellular component of Gene Ontology enrichment analysis of all proteins identified in the different tested conditions. Biological process, Molecular function and KEGG Gene Ontology analysis are shown on extended figure 3. **c**, Protein-Protein Interaction Network Exploration of the pooled proteins identified in the cellular component Gene Ontology enrichment analysis for potential PA interactors under different conditions using Cytoscape (STRING data base). Circles highlight clusters of proteins involved in interactions: red (H^+^ transport by V-ATPase), blue (SNAREs), yellow (mitochondrial activity), green (membrane coats).

Gene ontology analysis classifying proteins based on molecular function, biological process, and cellular component revealed specific enrichments in proteins associated with membrane organelles and trafficking pathways. Notably, we observed clear differences between PA species (Figure 6b and extended Figure 5). Furthermore, although potential partners of PA (18:0-18:1) were less abundant, they showed a more distinct pattern across the three cellular states tested, with a dramatic increase in response to stimulation (Figure 6b and extended Figure 5).

The potential PA partners inferred to the GO terms “cellular components” were pooled and STRING maps were generated to visualize the interaction networks (Figure 6c). Interestingly, CgA, known for its role in secretory granule biogenesis through its interaction with PA^33^, was found to interact with both synthetic PA **21a** and **21d** across all conditions (Fig. 6c, boxed in black). For PA (18:0-18:1) **21a** an increase in the number of potential partners was observed after stimulation, including proteins related to the vacuolar ATPase (V-ATPase), which has been implicated in PA formation near exocytotic site^45^ (Figure 4c). Additionally, SNARE-related proteins were detected specifically for PA-DDA1 (18:0-22:6) **21d** (Figure 6c) in the stimulated condition, consistent with previous observations that PA may interact with syntaxin-1A^46^. Of interest, adaptor and membrane coat-related proteins were also detected post-stimulation, possibly related to recycling of granular content by compensatory endocytosis. The newly identified potential partners could be direct or indirect interactors. Thus direct interaction remains to be probed directly using biochemical approaches. Furthermore, detailed bioinformatic analysis of these protein sequences will provide more information to define PA binding motifs^40^.

To our knowledge, this study represents the first example of GPL-protein interatomic, or “fishing” experiment performed with GPLs embedded in membranes from primary living cells under different physiological conditions. Among the many novel interactants identified, several interesting candidates are involved in lipid transfer, metabolism, and translation highlighting new potential avenues for research.

## Conclusion

This work provides access to patented versatile and clickable PA analogues, that preserve natural fatty acyl chains and the phosphate headgroup, thus confirming their potential for *in cellulo* applications^47^. Unlike currently commercially available PA analogues, modified on their fatty acyl chains, these synthetic analogues faithfully reproduce the biological behavior of natural lipids. This highlights importance of fatty acyl chain composition for biological studies involving PA and probably other GPLs. Finally, fishing experiments performed on primary cells not only confirmed known interactions but also revealed dozens of others unknown interactions, depending both on the fatty acyl chain and the cellular state. Overall, these novel synthetic GPL significantly broaden our knowledge of PA partners and functions, opening new avenues for exploring lipid-mediated cellular processes, lipid interactome and beyond.

## Fundings

This work was supported by grants from “Fondation pour la Recherche Médicale” (DEI20151234424) and “Agence Nationale pour la Recherche” (ANR-19-CE44-0019) to NV, PYR, SB and MM-H, “Fondation pour la Recherche Médicale” (FDT202304016618) to AW, French Proteomic Infrastructure (ProFI; ANR-10-INBS-08–03) to JMS. INSERM is providing the salary to NV and SG. TF and FL are co-supported by European Union and Région Normandie. This work has been partially supported by Université Rouen Normandie, INSA Rouen Normandie, Centre National de la Recherche Scientifique (CNRS), European Regional Development Fund (ERDF), Labex SynOrg (ANR-11-LABX-0029), Carnot Institute I2C, and the graduate school for research XL-Chem (ANR-18-EURE-0020 XL CHEM).

### Acknowledgements

We acknowledge the In Vitro Imaging Platform - NeuroPôle - Strasbourg (UAR3156), member of the national infrastructure France-BioImaging supported by the French National Research Agency (ANR-10-INBS-04). We acknowledge the municipal slaughterhouse of Haguenau (France) and the slaughterhouse of Bigard Formerie (France) for providing the bovine adrenal glands. We acknowledge the members of the Plateforme C2I Orga, specifically Laetitia Bailly, Emily Petit, Albert Marccual, Mathilde Lauzent and Françoise Ringot. SB thanks Dr. Antoine Joosten and Dr. Alexandre Haefelé for the scientific discussions about this project.

Université Rouen-Normandie, CNRS, INSA Rouen-Normandie and INSERM have issued a common patent (FR20220002640 20220324) based on this work. AS, AW, AC, LR, RF, PYR, MM-H, NV and SB are named as inventors on this patent. The other authors declare no competing interests. Data availability can be requested to the corresponding authors.

## Contributions

PYR, MM-H, NV and SB developed the initial concept. The synthesis of PA-N_3_ were performed by AS, AC, RF and BB. The synthesis of DIBAC was performed by LG, AS, SV, PJH and BB. The synthesis of DIBAC-C6-NBD was performed by AS and PJH. The synthesis of DIBAC-C6ATTO was performed by AS. The synthesis of PCL-DDA1 was performed by SV. The synthesis of PA-NBD analogues was performed by PJH, AC and AS. The synthesis of PA-ATTO analogues was performed by AS. The synthesis of PA-DDA1 were performed by SV and CS. The NMR study about transesterification resulting from the synthesis of the phospholipid model was performed by MS, LG, AP and SV. Purifications of final probes were performed by VF, VAP and PC. GUV experiments were performed by TF, LR and MC. Chromaffin cell cultures were managed by AW, ET, TF, FL and LJ. Live-cell imaging was conducted by TF, MC and MB. AW performed PA sensor recruitment experiments, exocytotic site labeling experiments, amperometry experiments and all subsequent analysis. AW and ET performed the fishing experiments. JMS performed the mass spectrometry analysis. CD, AW, SO, and NV performed the bioinformatic analysis. SB, NV, PYR and MM-H discussed the results and wrote the paper. AS, AW, TF, AC, SG, ET edited the paper.

