## Extended figures for "Designing new natural-mimetic phosphatidic acid: a versatile and innovative synthetic strategy for glycerophospholipid research"

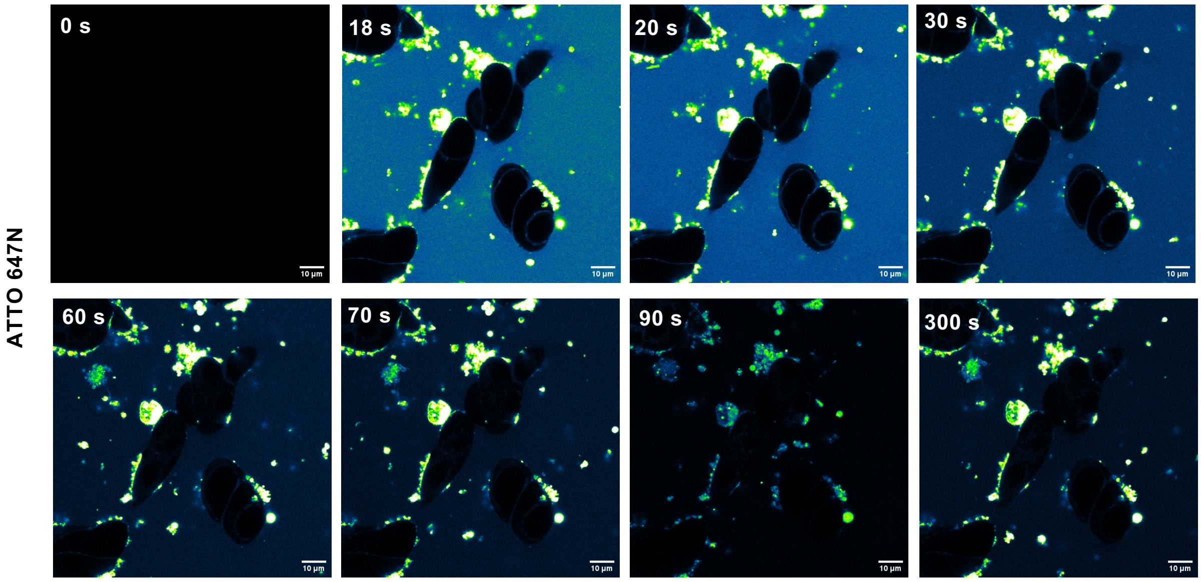


**Extended Figure 1: Live-cell imaging of ATTO 647N 3 µM incorporation in bovine chromaffin cells.** ATTO 647N has been added on cultured cells 15 s after the beginning of recording. Images have been false colored with ImageJ's Green Fire Blue LUT. Scale bar, 10 µm.

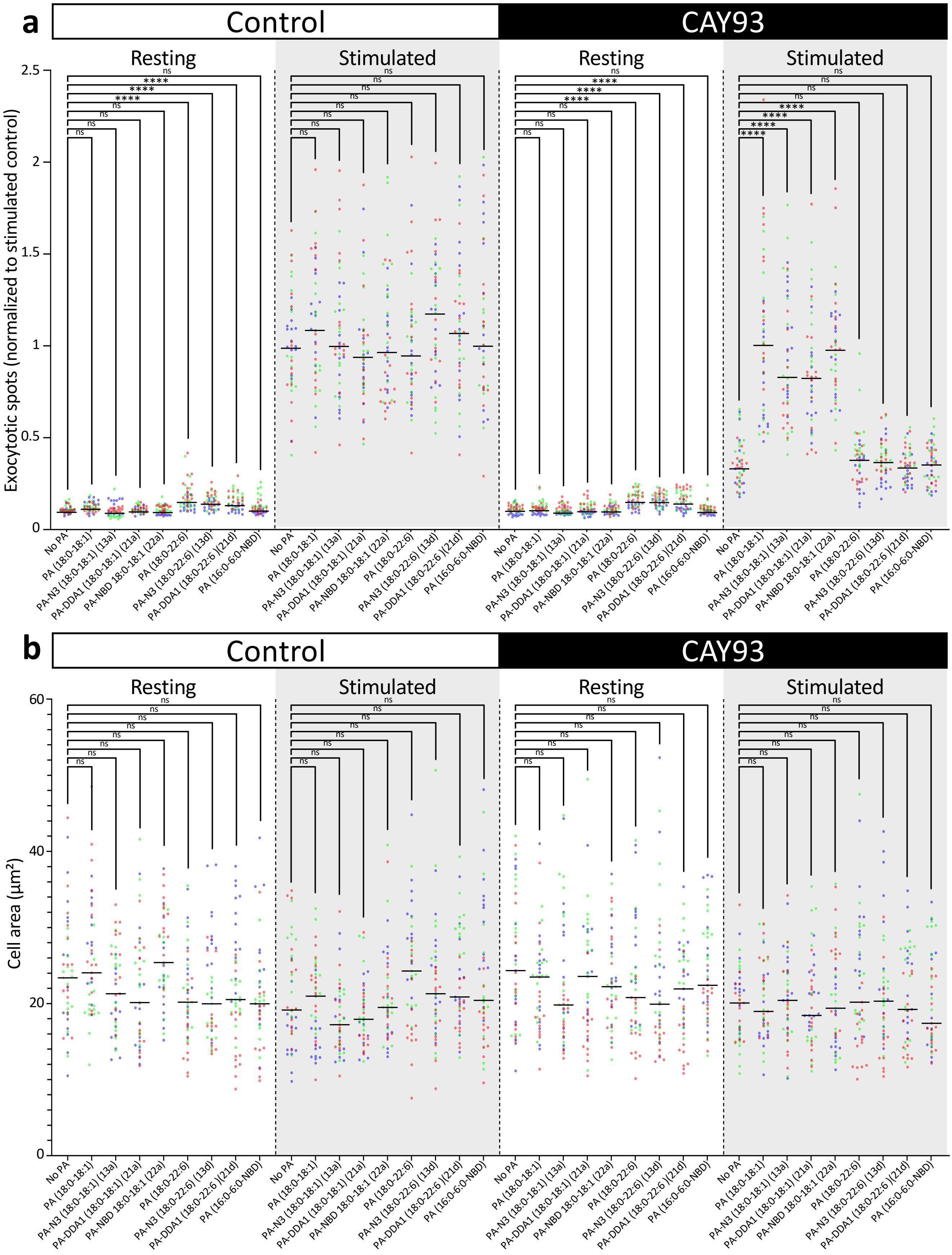
a
Control
CAY93
Resting
Stimulated
Exocytotic spots (normalized to stimulated control)
0
0.5
1
1.5
2
2.5
ns
****
No PA
PA (18:0-18:1)
PA-N3 (18:0-18:1) (13a)
PA-DDA1 (18:0-18:1) (21a)
PA-NBD 18:0-18:1 (22a)
PA (18:0-22:6)
PA-N3 (18:0-22:6) (13d)
PA-DDA1 (18:0-22:6 )(21d)
PA (16:0-6:0-NBD)
b
Control
CAY93
Resting
Stimulated
Cell area (μm²)
0
20
40
60
ns
No PA
PA (18:0-18:1)
PA-N3 (18:0-18:1) (13a)
PA-DDA1 (18:0-18:1) (21a)
PA-NBD 18:0-18:1 (22a)
PA (18:0-22:6)
PA-N3 (18:0-22:6) (13d)
PA-DDA1 (18:0-22:6 )(21d)
PA (16:0-6:0-NBD)

**Extended Figure 2: Effect of the different PA analogues on exocytosis analyzed by measuring cell surface exocytotic sites.** a, Quantification of peripheral exocytotic spot of bovine chromaffin cells incubated with DMSO (vehicle), 50 nM CAY93, and the indicated PA or PA analogues in resting (3 min Locke's solution) or stimulated (3 min, 59 mM $K^+$) condition. Mean peripheral exocytotic spot intensity has been normalized with the mean control condition of the individual experimental repeat. b, Quantification of the cellular area of each cell. For both graphs, data is represented as a median and each point is a measure from an individual cell. Symbols are color coded (red, blue or green) according to experimental repeats. ns: nonsignificant, ****: $p<0.0001$, Kruskall-Wallis followed by Dunn's multiple comparisons test vs control (N=45 per condition, 15 cells from 3 independent cultures).

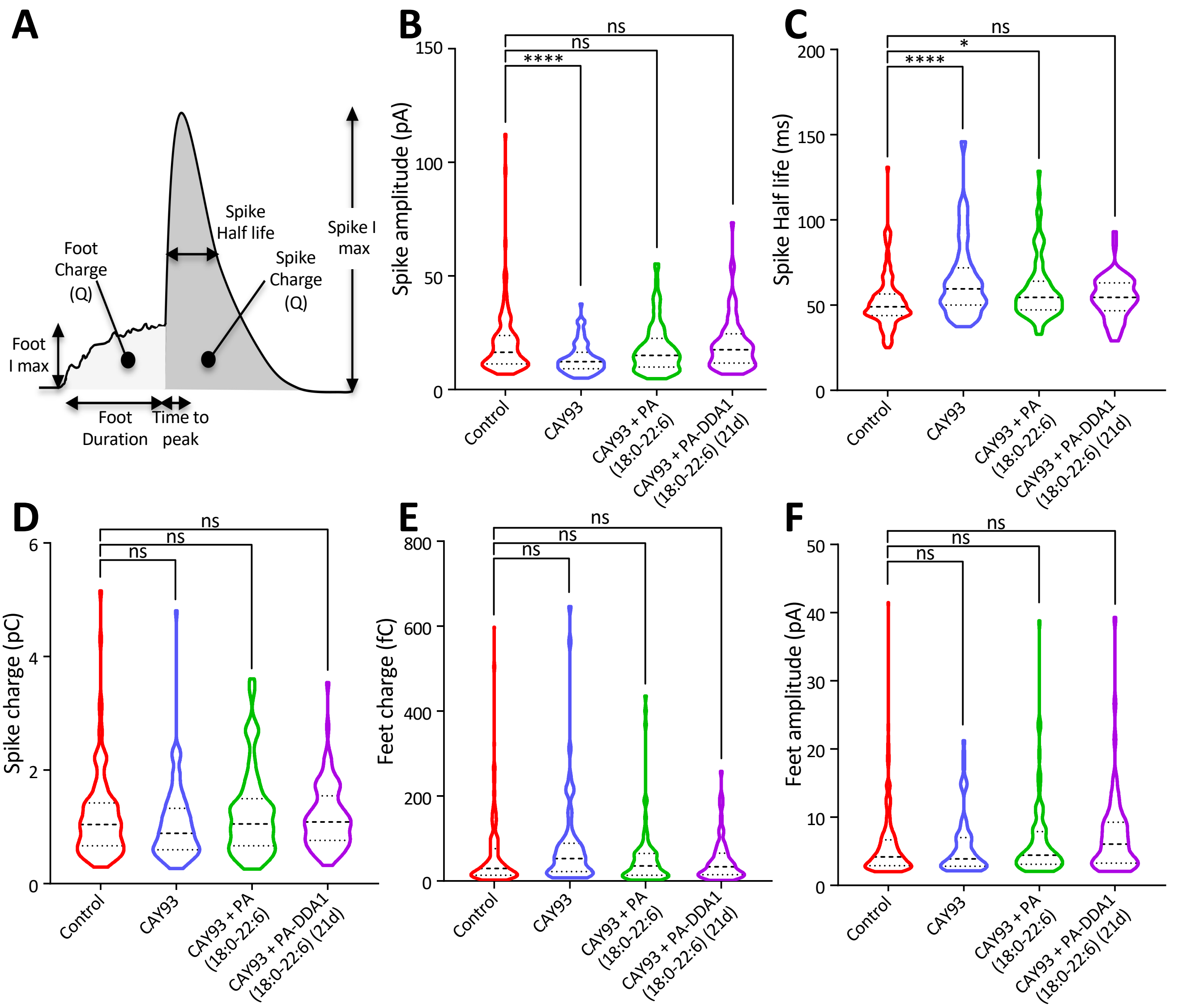


**Cell parameter**

**G**

| | Control | CAY93 | CAY93 + PA (18:0-22:6) | CAY93 + PA-DDA1 (18:0-22:6) (21d) |
|---|---|---|---|---|
| Analyzed cells | 115 | 91 | 86 | 82 |
| Total spikes | 11.99 ± 0.05 | 6.51 ± 0.04 (****) | 6.77 ± 0.04 (****) | 6.95 ± 0.04 (****) |
| Analyzable spikes | 9.49 ± 0.04 | 5.37 ± 0.03 (****) | 5.23 ± 0.03 (****) | 5.28 ± 0.03 (****) |
| Imax (pA) | 20.74 ± 0.08 | 13.76 ± 0.06 (****) | 18.10 ± 0.09 | 19.83 ± 0.10 |
| Charge (pC) | 1.19 ± 0.005 | 1.05 ± 0.005 | 1.24 ± 0.006 | 1.18 ± 0.005 |
| Half life (ms) | 52.16 ± 0.10 | 64.78 ± 0.18 (****) | 59.40 ± 0.15 (*) | 54.66 ± 0.12 |
| Time to peak (ms) | 25.04 ± 0.05 | 35.31 ± 0.11 (****) | 28.70 ± 0.08 | 26.16 ± 0.07 |

**Individual spike parameter**

| | Control | CAY93 | CAY93 + PA (18:0-22:6) | CAY93 + PA-DDA1 (18:0-22:6) (21d) |
|---|---|---|---|---|
| Analyzed spikes | 1091 | 489 | 452 | 433 |
| Imax (pA) | 20.52 ± 0.01 | 14.92 ± 0.02 (****) | 20.87 ± 0.03 | 22.52 ± 0.03 (*) |
| Charge (pC) | 1.14 ± 0.001 | 1.08 ± 0.001 | 1.33 ± 0.002 (***) | 1.29 ± 0.002 (***) |
| Half life (ms) | 50.18 ± 0.01 | 62.72 ± 0.04 (****) | 55.77 ± 0.03 (****) | 53.26 ± 0.03 (****) |
| Time to peak (ms) | 24.15 ± 0.01 | 33.70 ± 0.02 (****) | 26.71 ± 0.02 (***) | 25.88 ± 0.02 (**) |

**Feet parameter**

| | Control | CAY93 | CAY93 + PA (18:0-22:6) | CAY93 + PA-DDA1 (18:0-22:6) (21d) |
|---|---|---|---|---|
| Analyzed feet | 208 | 64 | 86 | 66 |
| Imax (pA) | 5.80 ± 0.01 | 5.74 ± 0.05 | 6.37 ± 0.04 | 7.40 ± 0.06 |
| Charge (fC) | 60.74 ± 0.26 | 86.22 ± 1.08 | 55.29 ± 0.51 | 49.07 ± 0.56 |
| Duration (ms) | 25.34 ± 0.07 | 37.35 ± 0.29 (****) | 22.77 ± 0.12 | 20.54 ± 0.15 |

**Extended Figure 3: Effect of the different PA analogues on exocytosis analyzed by carbon fiber amperometry.** a, Schematic representation of the different individual spike parameters that have been quantified. Violin plots representing respectively the spike amplitude b, half life c, charge d, feet charge e, and amplitude f, of the amperometry experiment (Bar in the plot represent form top to bottom, 3$^{rd}$ quartile, median and 1$^{st}$ quartile). g, Table recapitulating the different parameters that have been analyzed in the amperometry experiment. ns: nonsignificant, *: p<0.05, **: p<0.01, ***: p<0.001, ****: p<0.0001, Kruskall-Wallis followed by Dunn's multiple comparisons test compared to control.

| Interactant | Function | PA-DDA1 (18:0-18:1) (21a) | PA-DDA1 (18:0-22:6) (21d) | Reference |
|---|---|---|---|---|
| ANXA2 | Annexin, $Ca^{2+}$-dependent lipid binding | + | + | 1 |
| ANXA3 | Annexin, $Ca^{2+}$-dependent lipid binding | | + | 2 |
| ARF1 | GTPase | + | | 3 |
| BANF1 | Nuclear assembly | + | | 1 |
| CERT | Lipid transport protein | + | | 4 |
| CGA | Secreted peptide and vesicle budding | + | + | 5 |
| CLTC | Coat protein | + | + | 1 |
| COPB1 | Coat protein | + | + | 3 |
| DDX3Y | Protein translation | | + | 1 |
| DNM1L | Membrane fission | | + | 1 |
| EIF4A2 | Translation initiator | + | | 1 |
| EIF4A3 | Translation initiator | | + | 1 |
| FAU | Ribosomal protein | | + | 1 |
| HNRNPR | Nuclear ribonucleoprotein | | + | 1 |
| KIF5B | Kinesin, vesicular transport | | + | 3 |
| MCCC2 | Methylcrotonyl-CoA carboxylase subunit | + | | 1 |
| MYH10 | Myosin, vesicular transport | + | | 1 |
| MYH9 | Myosin, vesicular transport | | + | 1 |
| NAPA | SNARE associated protein | | + | 6 |
| NSF | SNARE associated protein | + | + | 3,6 |
| PARK7 | Parkinsonism associated deglycase | | + | 1 |
| PCCA | Propionyl-CoA Carboxylase Subunit | + | | 1 |
| PITPNA | Lipid transport protein | + | + | 7 |
| PITPNB | Lipid transport protein | + | + | 7 |
| PP1 | Protein phosphatase 1 catalytic subunit | + | + | 8 |
| PPP1CC | Protein Phosphatase 1 Catalytic Subunit | + | + | 8,9 |
| RAC1 | GTPase | + | | 10,11 |
| RBM14 | RNA binding protein | + | | 1 |
| RHOG | GTPase | | + | 2 |
| RPL10L | Ribosomal protein | | + | 1 |
| RPL15 | Ribosomal protein | + | | 1 |
| RPL21 | Ribosomal protein | + | | 1 |
| RPL24 | Ribosomal protein | + | | 1 |
| RPL3 | Ribosomal protein | + | + | 1 |
| RPL35 | Ribosomal protein | | + | 1 |
| RPL37A | Ribosomal protein | | + | 1 |
| RPL9 | Ribosomal protein | + | + | 1 |
| RPLP0 | Ribosomal protein | + | | 1 |
| RPS11 | Ribosomal protein | + | | 1 |
| RPS16 | Ribosomal protein | + | + | 1 |
| RPS24 | Ribosomal protein | + | | 1 |
| RPS26 | Ribosomal protein | | + | 1 |
| RPS27 | Ribosomal protein | + | | 1 |
| RPS9 | Ribosomal protein | + | | 1 |
| SNCA | α-synuclein | | + | 12 |
| SRSF7 | Protein splicing factor | + | | 1 |
| STAT3 | Transcription factor | + | | 1 |
| STX1A | SNARE protein | + | + | 13 |
| TRA2B | Protein splicing factor | + | + | 1 |
| TRIM59 | E3 ubiquitin ligase | | + | 1 |
| TRMT112 | tRNA methylation | + | | 1 |
| TUBA1C | Tubulin | + | + | 1 |
| UBA52 | Ribosomal protein | + | | 1 |
| VAPA | Lipid transport protein | + | + | 1,14 |
| VAPB | Lipid transport protein | + | + | 1 |

**Extended Figure 4: List of the synthetic PA 21a and 21d partners and their corresponding function identified in our screen and found in previous studies using natural PA with the corresponding references.**

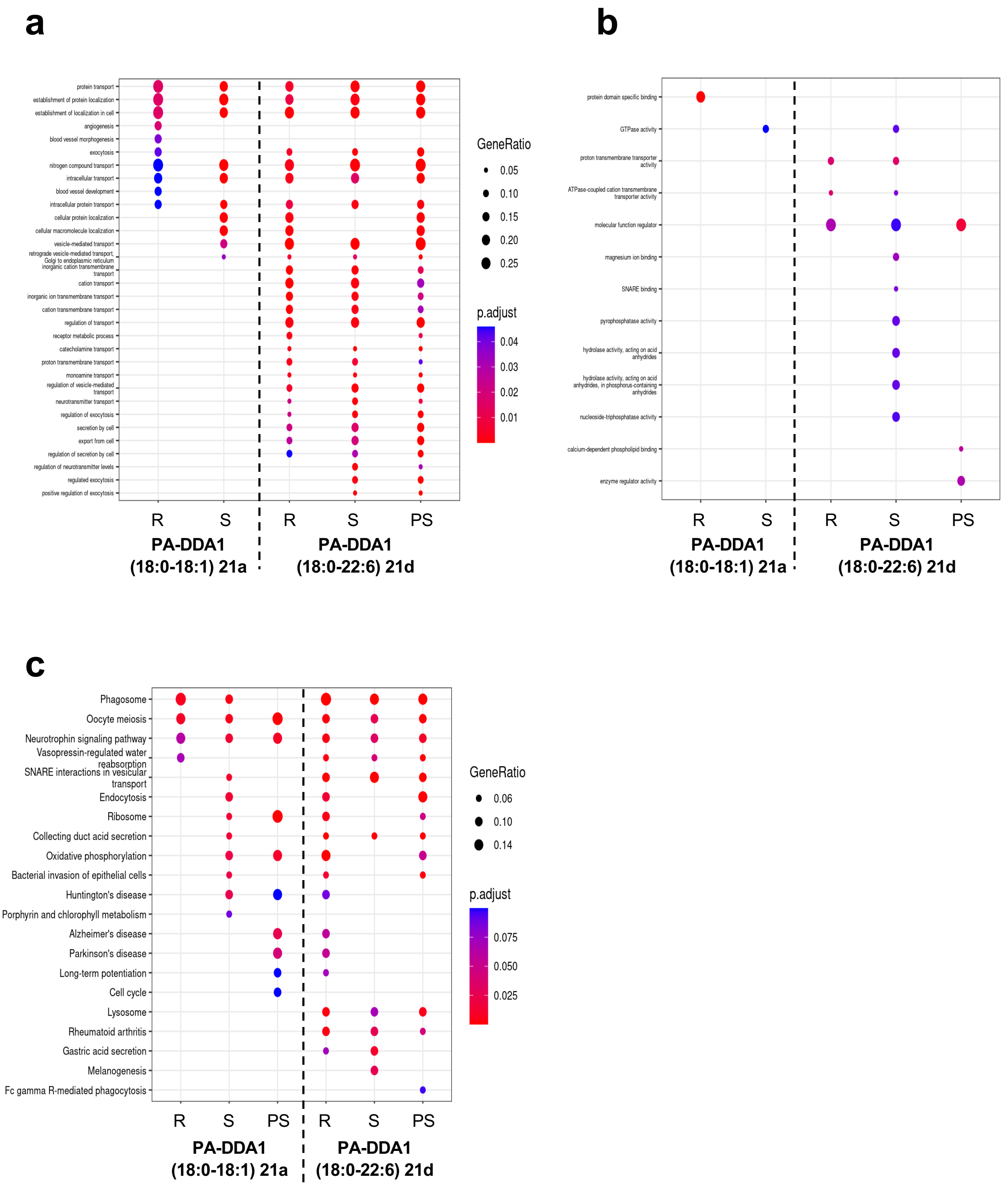


**Extended Figure 5: Functional enrichment analysis of PA fishing experiments.** Gene ontology biological processes enrichment analysis (**a**), gene ontology molecular functions enrichment analysis (**b**), and KEGG pathways analysis (**c**) of the proteins identified in the fishing experiments performed in bovine chromaffin cells for PA-DDA1 (18:0-18:1) **21a** and PA-DDA1 (18:0-22:6) **21d** in resting (R), stimulated (S), or after stimulation (PS).
